# Epigenetic upregulation of carotid body angiotensin signaling increases blood pressure

**DOI:** 10.1101/2024.05.10.593589

**Authors:** Fengli Zhu, Zhuqing Wang, Kayla Davis, Hayden McSwiggin, Jekaterina Zyuzin, Jie Liu, Wei Yan, Virender K. Rehan, Nicholas Jendzjowsky

## Abstract

Epigenetic changes can be shaped by a wide array of environmental cues as well as maternal health and behaviors. One of the most detrimental behaviors to the developing fetus is nicotine exposure. Perinatal nicotine exposure remains a significant risk factor for cardiovascular health and in particular, hypertension. Increased basal carotid body activity and excitation are significant contributors to hypertension. This study investigated the epigenetic changes to carotid body activity induced by perinatal nicotine exposure resulting in carotid body-mediated hypertension. Using a rodent model of perinatal nicotine exposure, we show that angiotensin II type 1 receptor is upregulated in the carotid bodies of nicotine-exposed offspring. These changes were attributed to an upregulation of genetic promotion as DNA methylation of angiotensin II type 1 receptor occurred within intron regions, exemplifying an upregulation of genetic transcription for these genes. Nicotine increased angiotensin signaling *in vitro*. Carotid body reactivity to angiotensin was increased in perinatal nicotine-exposed offspring compared to control offspring. Further, carotid body denervation reduced arterial pressure as a result of suppressed efferent sympathetic activity in perinatal nicotine-exposed offspring. Our data demonstrate that perinatal nicotine exposure adversely affects carotid body afferent sensing, which augments efferent sympathetic activity to increase vasoconstrictor signaling and induce hypertension. Targeting angiotensin signaling in the carotid bodies may provide a way to alleviate hypertension acquired by adverse maternal uterine environments in general and perinatal nicotine exposure in particular.

## Introduction

Adverse developmental programming is directly related to cardiovascular disease in later life. The epigenetic modifications that occur in response to intra-uterine alterations have lasting effects and are known to alter cardiovascular structure and function in response to several environmental factors and maternal behaviors^1–5^. One of the most detrimental maternal behaviors during the perinatal period, which can deleteriously predispose offspring to cardiovascular diseases, is nicotine exposure, typically in the form of tobacco product use^6^.

Perinatal tobacco smoke/nicotine exposure has been shown to augment blood pressure and increase the risk of hypertension in later life^2,6,7^. Arteriolar wall thickening and increased vasoconstrictor signaling are thought to be the primary causes of perinatal smoke/nicotine exposure-induced hypertension^8–10^. Nicotine stimulates the renin-angiotensin-aldosterone system^11^ and increases both angiotensin-converting enzyme activity^12,13^ as well as angiotensin II type 1 receptor (AgtR1) expression and activity^12–14^ within the aorta^15–17^ and mesenteric arterioles^16,18^. Such arteriolar reactivity would likely implement a hypertensive state.

Aberrant sympathetic nerve activity is significantly tied to cardiovascular disease and a prime mediator of hypertension in a multitude of cardiometabolic diseases^19,20^. Carotid bodies are polymodal sensors that regulate sympathetic nervous activity to control vasoconstriction to regulate tissue oxygen/nutrient delivery^21,22^. Indeed, hypertension appears to be critically dependent on tonic input from carotid bodies to augment sympathetic tone^19,20^. Carotid body activity of nicotine-exposed offspring is not suppressed by hyperoxia, which typically quiesces carotid body activity, suggesting an augmented basal sensitization^23,24^. Given their primary role in stimulating arteriolar vasoconstriction^19,20^ and their susceptibility to nicotine exposure^23,24^, the carotid bodies are likely susceptible to epigenetic modulation because of developmental programming^25–27^, which would lead to an elevation of tonic activity. Therefore, we investigated the hypothesis that carotid bodies are susceptible to epigenetic hypersensitization which results in increased blood pressure in perinatal nicotine-exposed offspring. To test our hypothesis, we engaged modern unbiased sequencing of RNA and DNA methylation of carotid bodies, tested signaling pathways *in vitro* and *in vivo,* and assessed the necessity of carotid bodies for heightened arterial pressure in perinatal nicotine-exposed pups.

## Methods

### Data availability

RNA sequencing and DNA bisulfite sequencing data have been deposited and can be found in the NIH SRA (Bioproject ID: PRJNA1106547). All other data are found in the supporting data and supplementary files and can be further made available upon request to the corresponding author.

### Animal models, care and ethical standards

First-time pregnant Sprague-Dawley (SD) rat dams were purchased from Charles River (Hollister, CA) at gestational day 3. SD mothers were subject to subcutaneous nicotine injections (2mg/kg, Millipore Sigma, PHR2532) from gestational day 7 to post-natal day (PND) 21^28–30^. Control mothers received PBS (vehicle). Animals were housed with controlled temperature (21 ± 2 °C), humidity (55 ± 10%) and 12h light-dark cycle. Pups were weaned on post-natal day (PND) 21 and housed in pairs. Animals had unlimited access to food and water. Experiments on pups took place from PND56-80. All experimental procedures were approved by The Lundquist Institute for Biomedical Innovation at Harbor UCLA Medical Center IACUC board, protocol # 31659.

Assignment of control or nicotine to mothers was block randomized. Unbiased transcriptomic and genomic sequencing was based on previous data acquired by neuronal sequencing results. *In vitro* and *in vivo* power calculations were based on an estimated effect size of 50% difference with assumed unified standard deviation across groups.

### PC12 culture

PC12 cells were cultured in RPMI 1640 (Gibco # 11875093) supplemented with 10% Horse Serum (Gibco Cat. #26050-088) and 5% fetal bovine serum (Gibco Cat. #F2442), 0.4µg/ml dexamethasone (Millipore Sigma Cat. # D4902) and 1% penicillin/streptomycin (Millipore Sigma Cat. # P0781). Cells were seeded at 3×10^5^ cells in 1ml of media per well (24 well plate, qPCR) or 2500 cells in 150µl of media per well (96 well plate calcium imaging).

### Next-generation RNA sequencing

On PND 56, animals were sedated (5% isoflurane, balance O2), exsanguinated and carotid bodies gross dissected and placed in a sylgard-coated dissecting dish filled with Ham’s F12 media. Carotid bodies were freed from the carotid bifurcation under a dissecting microscope. Five carotid body pairs (n=10 carotid bodies from 5 rats) from each group and sex (n=5 rats per group, per sex) were lysed and RNA was isolated using the easy-spin Total RNA extraction kit (Boca scientific, Cat. # 17221). RNA purity was tested with Nanodrop spectrophotometer (Thermofisher Cat. # ND2000USCAN) and deemed to be of appropriate quality when A260/280 was between 1.99 and 2.10. RNA sequencing was then conducted by an unbiased commercial vendor (Novogene corporation, Sacramento CA), where further RNA purity was analyzed (RIN >7) and downstream library construction took place.

### Whole genome DNA-bisulfite sequencing

Carotid bodies were dissected as above. Five carotid body pairs (n=10 carotid bodies from 5 rats) from each group and sex (n=5 rats per group, per sex) were lysed and DNA was isolated using the QIAamp UCP DNA Micro Kit (Qiagen, Cat. # 56204). DNA purity was deemed sufficient with A260/280 between 1.75-1.85. Bisulfite conversion and downstream whole-genome DNA-bisulfite sequencing were conducted by (Novogene Corporation, Sacramento, CA).

### RNA qPCR

For carotid bodies, an additional group of n=3 samples (n=5 rats per sample) from each group was harvested as above, snap frozen and stored until analysis. Brain and kidneys were harvested from each group of pups (one tissue per pup was deemed as n=1). For PC12 cells were washed in PBS and then immediately lysed in lysis buffer and converted into cDNA. N=3 wells were analyzed for each condition for each probe and five separate experiments were run. RNA was converted to cDNA using the Tetro cDNA synthesis kit (Meridian life sciences, Cat. # Bio65043). qPCR was conducted with Taqman specific probes. AgtR1a (Thermofisher Rn02758772_s1), AgtR2 (Thermofisher Rn00560677_s1), angiotensinogen (Thermofisher RN00593114_m1), angiotensin-converting enzyme (Thermofisher Rn00561094_m1), renin (Thermofisher Rn02586313_m1), TH (Thermofisher Rn00562500_m1), endothelial PAS domain-containing protein 1/ hypoxia inducible factor 2 alpha (EPAS1/Hif2α, Rn00576515_m1) and TRPV1 (Thermofisher Rn00583117_m1). All experimental genes were evaluated in reference to hypoxanthine-guanine phosphoribosyltransferase (HPRT Thermofisher Rn01527840_m1). The qPCR reaction was completed using Taqman fast advanced master mix enzyme (Thermofisher Cat. # 4444557) and the QuantStudio 3 thermocycler (Thermofisher Cat. #A28567). Delta CT was calculated in reference to HPRT, and comparisons between groups were made using an unpaired two-sided t-test.

### Methylated DNA immunoprecipitation-qPCR (MeDIP-qPCR)

DNA was isolated from carotid bodies (n=4 per group, each n is comprised of carotid bodies from 5 rats), brains and kidneys (n=4 rats per group, one tissue is n=1) and PC12 cells (1 well is n=1). Carotid body and PC12 DNA was isolated using the QIAamp UCP DNA Micro Kit (Qiagen, Cat. # 56204). Brain and kidney DNA was isolated using the Puregene Tissue kit (Qiagen, Cat. # 158063). MeDIP was performed using the Methylated DNA immunoprecipitation kit (Cat.# ab117133, Abcam) following the procedures described previously^31^. Briefly, 100μL of the antibody buffer and 1μL anti-5-methylcytosine or reference IgG antibody were added into wells and incubated at room temperature for 1 hour. During the incubation, 1μg of genomic DNA was sheared between 200 and 1000 bp in the reaction buffer using the Covaris M220 ultrasonicator, followed by denaturation at 95 °C for 2 minutes, then placed on ice until use. An aliquot of 5 μL of the denatured DNA was saved as the input DNA. The strip wells bound with antibodies were washed with 150μL of the antibody buffer and 150μL of the wash buffer once, followed by incubation with the sheared DNA at room temperature for 2 hours. The antibody-enriched DNA was eluted in the DNA release buffer containing proteinase K followed by DNA purification. qPCR was performed using Fast SYBR™ Green Master Mix (Cat.# 4385612, Thermo Fisher Scientific) to identify DNA methylation levels. Primer sequence-AgtR1: Forward TACCTAAACATAGTAAAAGCCAAACACA; AgtR1: Reverse TATCCTGTTGATCTCTTTTTGTTGTCTG. ΔCt was calculated as Methylated AgtR1 DNA – Total AgtR1 DNA and was used to plot the data^31,32^. Comparisons between groups were made using an unpaired two-sided t-test.

### Immunohistochemistry

Carotid bodies were gross dissected as above. Then they were cleaned from connective tissue but left attached to the surrounding carotid artery bifurcation as previously^33^. Tissues were fixed in 4% paraformaldehyde for 1-2 hours at 4°C, then cryopreserved in 30% sucrose overnight at 4°C, dried embedded in optimal cutting temperature compound (OCT) and cut at 14µm sections on a cryostat. Sections were then washed (PBS), permeabilized (Triton x 20%) blocked with 10% goat serum (Abcam, Cat. # ab7481), and incubated with antibodies for tyrosine hydroxylase (TH, Millipore Sigma ZMS1033, clone 20/40/15), AgtR1 (Millipore Sigma AB15552, polyclonal) and TRPV1 (Alomone Labs ACC-030, polyconal) for 24hours at 4°C and subsequently stained with appropriate secondaries (Jackson Laboratories Goat-anti rabbit Cy3 111-165-003, Cy5 111-005-003 and Goat-anti mouse 488 115-545-003) for 90 minutes and then counterstained with DAPI (Millipore Sigma MBD0015) and coverslipped with prolong diamond antifade mounting media (Thermofisher #P36970). Slides were imaged using the Leica Thunder imaging acquisition system.

### Calcium imaging

PC12 cells were separated into media (control), stimulated with nicotine (Sigma Cat. # 6019-06-3, 50µg/ml), angiotensin II (Tocris Cat. # 1158, 5µM), nicotine + angiotensin II, angiotensin II + losartan (Tocris Cat. # 3798, 3µM), nicotine + angiotensin II + losartan, angiotensin II + AMG9810 (TRPV1 antagonist Tocris Cat. # 2316, 10µM), nicotine + angiotensin II + AMG9810, or nicotine + angiotensin II + losartan + AMG9810, for 1 hour and then incubated with Calcium Orange AM methyl ester (Invitrogen Cat. # C3015) + 0.01% Pluronic acid (Invitrogen Cat. #F-127) for 20 minutes, washed in media and then imaged in the Incucyte imaging system (Sartorius) using the Neural imaging parameters (Ex: 549λm/ Em: 576λm, 60s) for 1 hour. The frequency of methyl ester excitation bursts was averaged for each well to express a single value. Data were analyzed with One-way ANOVA using the Holm-Sidak post-hoc test.

### In vivo carotid body reactivity

Rats from control or nicotine groups (as above) were tested on PND 56-58. Rats were anesthetized with 5% isoflurane (balance room air) using the Kent Scientific Somnosuite anesthesia system. The femoral artery and vein were cannulated, and the arterial line was instrumented to a pressure transducer to measure continuous arterial pressure (AD instruments MLT0670) using the PowerLab data acquisition console (AD instruments PL3516/P). The vein was then connected to a syringe pump (Fisherscientific Cat. # 14831200) to induce a 15mg/kg/min infusion of alfaxan (Victor Medical Cat. # 1629020)^34,35^. Then the jugular vein was cannulated to deliver bolus injections of Angiotensin II (Tocris Cat. # 1158 1mg/kg) and sodium cyanide (NaCN, Fisherscientific Cat. # AA1213722, 200µg/kg). These injections were repeated following bilateral carotid body denervation. Briefly, the neck area was shaved, and a midline incision exposed the carotid artery. Then glands were pushed aside, and the carotid bodies were identified within the carotid bifurcation, underneath the occipital artery. The carotid sinus nerve was confirmed as it was traced to the glossopharyngeal nerve, then denervated, as previous^34^. Mean arterial pressure was averaged at the 60s preceding each bolus injection and a 60s average during the peak reactive response to each bolus injection. Data were analyzed by Two-way ANOVA (Group x Carotid body intact vs. denervated) for Angiotensin II or NaCN injection. Holm-Sidak post-hoc test determined group differences.

### Serum angiotensin concentration

Blood was collected after induction of IV anesthetic but before any drug infusion from the above experiment from the femoral vein. Blood was left to coagulate for 20 minutes on ice then spun at 4,000rpm for 10 minutes at 4°C. Serum was drawn, aliquoted, snap-frozen, and stored at -80°C until analysis. Angiotensin ELISA (Raybiotech #EIAR-ANGII-1) was used per manufacturer’s instructions. Groups (Control pups vs. perinatal nicotine-exposed pups) were analyzed with two-sided unpaired t-test.

### Arterial pressure assessment between groups

Rats from control or perinatal nicotine-exposed mothers were handled daily following weaning at PND 21 and habituated to the tail plethysmography system (IITC Lifesciences). Three-minute averages of tail cuff-measured blood pressure was averaged to attain a single value for each animal. Data were compared using an unpaired two-sided t-test.

### Carotid body-mediated changes in arterial pressure

Perinatal nicotine-exposed rats were prepared for telemetric implant between PND 56-60. Rats were anesthetized with 5% isoflurane (balance O_2_) as induction and maintained on a surgical plane with 2-3% isoflurane. The abdominal area was isolated, shaved, and treated with 3x betadyne and 3x 70% alcohol. A midline incision was made, and the descending aorta below the renal arteries and renal nerves were isolated. The Kaha telemetry dual sympathetic nerve activity /pressure (TRM56SP, AD Instruments) were instrumented around the renal nerves using KwikGard dental impression material to isolate electrodes for renal sympathetic nerve activity (RSNA) recording. The pressure transducer was inserted into the aorta as per manufacturer’s instructions to measure continuous pressure measurement. The surgical area was sewn in layers, and the skin was secured with surgical staples. Rats were given buprenorphine (0.03mg/kg SQ) 2x daily for three days and allowed to recover for a total of 5 days before commencing recording. Pressure and RSNA were recorded for 1 week; then rats were anesthetized as above; the neck area was isolated, shaved, and treated with 3x betadine and 3x 70% alcohol. A midline incision was made on the neck, and the carotid bodies were identified behind the occipital arteries. The carotid sinus nerve was severed bilaterally, and the skin was sewn with subcuticular sutures. Buprenorphine was delivered as above, and recording commenced for another week following recovery. Arterial pressure waveforms delineated heart rate, systolic, diastolic, mean arterial pressure, pulse pressure, and respiratory rate. RSNA was filtered, rectified, and integrated using LabChart 8. Data were binned into daily averages and analyzed pre-post with a two-sided paired t-test.

### Statistics

Specific statistical tests are described for each experiment above. Data were averaged and analyzed for each experiment as above. Data were analyzed using Graphpad v10.

## Results

### Epigenetic changes in the carotid body in response to perinatal nicotine exposure

To investigate possible genetic changes expounded by adverse maternal behaviors, we implemented a standardized model of perinatal-nicotine exposure^28–30^. Given that the carotid bodies are prime drivers of hypertension in self-actualized cardiometabolic disease states^22,36^, we conducted an unbiased transcriptomic screen of carotid bodies harvested from perinatal nicotine-exposed pups on PND 56, 5 weeks after weaning and the final dose of nicotine delivered to the mother. We pooled together carotid bodies from 5 pups in each group of nicotine-exposed and saline control pups (Figure 1a). Samples were analyzed by sex, but no sex difference was found. A total of 922 genes were differentially expressed between conditions, with 685 up-regulated, 237 down-regulated, and 19196 unchanged transcripts (Figure 1b, c). AgtR1a was significantly upregulated in nicotine-exposed offspring compared to control (0.69 Log2 change, p_adj_=0.04). Angiotensin II type 2 receptor (AgtR2) was downregulated by perinatal nicotine exposure (Log2FC=-3.246, p_adj_ =6.65×10^-12^). No differences in angiotensinogen (Log2FC=0.233, p_adj_ =0.693), angiotensin-converting enzyme (Log2FC=-0.071, p_adj_=0.871), or renin (Log2FC=-2.248, p_adj_=0.645) were demonstrated (Supplementary data 2). Gene set enrichment analysis (GSEA) was performed on the differentially expressed gene, revealing a vast number of highly significant GO pathways, including GO:0004930 (normalized enrichment score =1.539; Figure1d, Supplementary data 1), which included AgtR1a.

**Figure 1.**
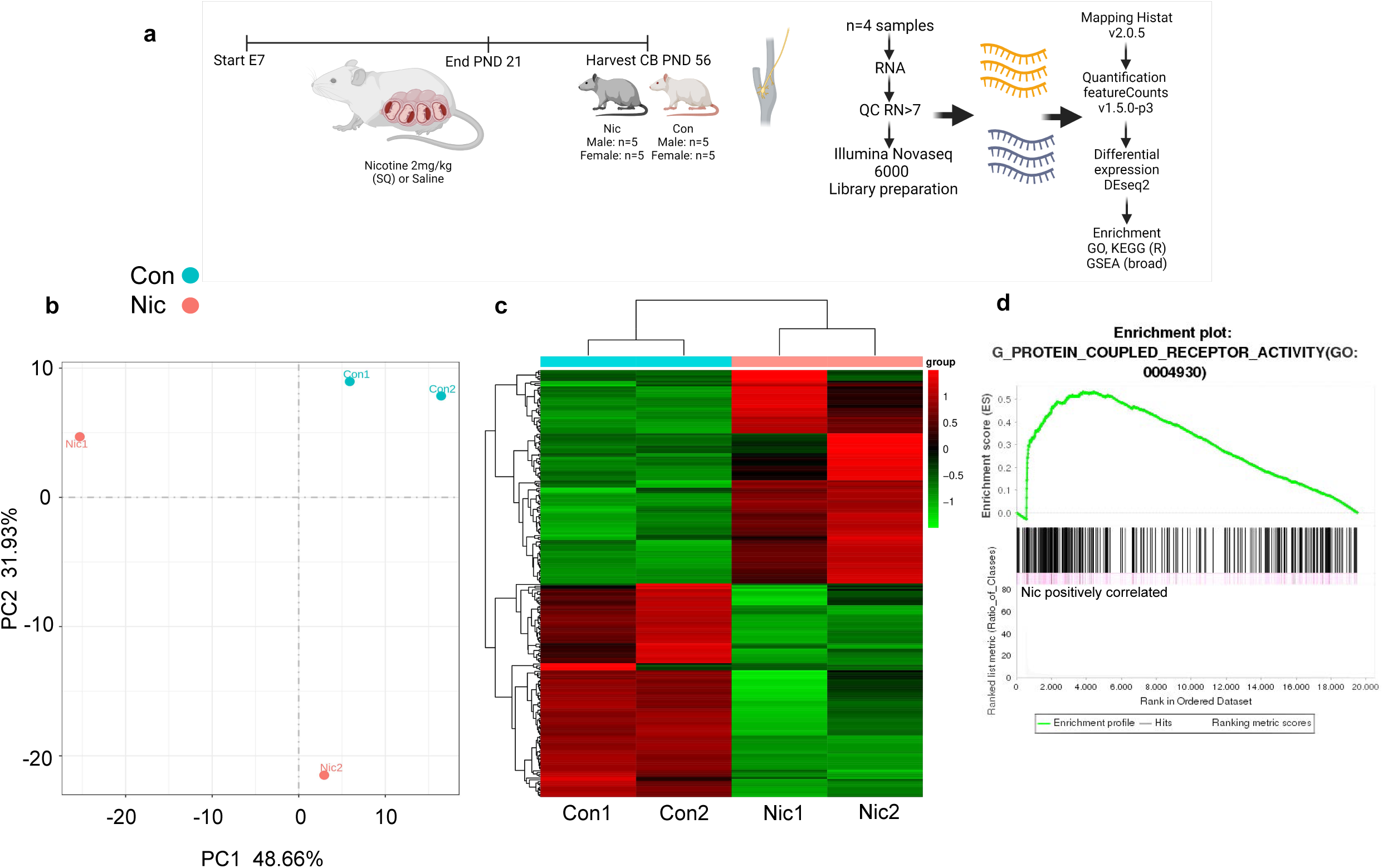
Perinatal nicotine exposure alters RNA transcripts associated with angiotensin signaling in the carotid body. **a)** Transcriptomic study design. Nicotine or saline was delivered to the mother on estrous day 7 (E7) to post-natal day (PND) 21. Bilateral carotid body (CB) samples were micro-dissected from perinatal nicotine or saline (control) exposed rats on PND 56. (n=5 rats per group-tissues were pooled to yield sufficient RNA for analysis) Two male and Two female groups per condition. **b)** Principal component analysis (PCA) plot showing distinct separation between nicotine and control groups. PCA1 was group and PCA2 was sex. **c)** Heatmap showing overall differences in RNA distribution. **d)** Gene set enrichment analysis (GSEA) plot for G-protein-coupled receptor activity (0004930) where AgtR1a was significantly increased in nicotine-exposed offspring compared to controls.

To validate these findings, we conducted immunohistochemistry and qPCR with taqman probes to identify possible upregulation of potential signaling pathways in the carotid bodies between perinatal nicotine exposed and control pups. We show that in relation to HPRT, AgtR1a was significantly upregulated in perinatal nicotine-exposed offspring compared to control. AgtR2 and TRPV1 transcription was not significantly increased (Supplementary figure 1).

To test whether changes in angiotensin signaling occurred outside the carotid bodies, we tested whether RNA transcripts of AgtR1a, AgtR2, angiotensinogen, angiotensin-converting enzyme, or renin were altered in the kidney and brains of perinatal nicotine exposed pups compared to controls. We observed that angiotensinogen and angiotensin-converting enzymes were significantly upregulated in the brains and kidneys of perinatal nicotine-exposed rats compared to the control, and renin was downregulated in the kidneys of perinatal nicotine-exposed rats compared to the control with no change in the brain (Supplementary figure 2). In contrast, both AgtR1a and AgtR2 were unchanged in the brains and kidneys of perinatal nicotine-exposed rats compared to control rats (Supplementary figure 2).

Given that our goal was to elucidate possible epigenetic differences responsible for mediating a neurogenic form of hypertension, we probed further and conducted whole genome DNA bisulfite sequencing^37^ of the carotid bodies from perinatal nicotine-exposed and control pups on PND 56 (Figure 2a). Samples were analyzed by sex, but no sex difference was found. Mapping the whole genome of the carotid bodies between groups revealed that the predominant changes to DNA methylation occurred in the promoter (suppression) and intron/exon regions (enhancement, Figure 2b). Methylation was distributed throughout the genome where a differential set of Genes were altered predominantly on the cytosine-guanine (CG) and cytosine-arginine/threonine/guanine-guanine (CHG) regions (Figure 2b). GO analyses of whole genes (Figure 2d) and promoter regions (Figure 2e) revealed similar GO terms as with transcriptomic changes. Importantly, a significant difference in methylation AgtR1a between groups was demonstrated with a difference of 0.448319 between perinatal nicotine exposed and control rats (Supplementary data 3). AgtR1 was orthogonally confirmed with immunoprecipitation of methylated DNA compared to total DNA amplifying AgtR1 to assess global methylation (Figure 2c). AgtR1 DNA was not significantly methylated in brains but was increased in the kidneys of perinatal nicotine-exposed rats compared to controls (Supplementary figure 3). Given that these methylation changes occurred in regions associated with the upregulation of angiotensin GPCR signaling, we wanted to investigate the functional outcomes of this pathway further. The combined transcriptomic changes and DNA methylation of the whole genome point to an upregulation of angiotensin signaling within the carotid bodies of perinatal nicotine-exposed offspring.

**Figure 2.**
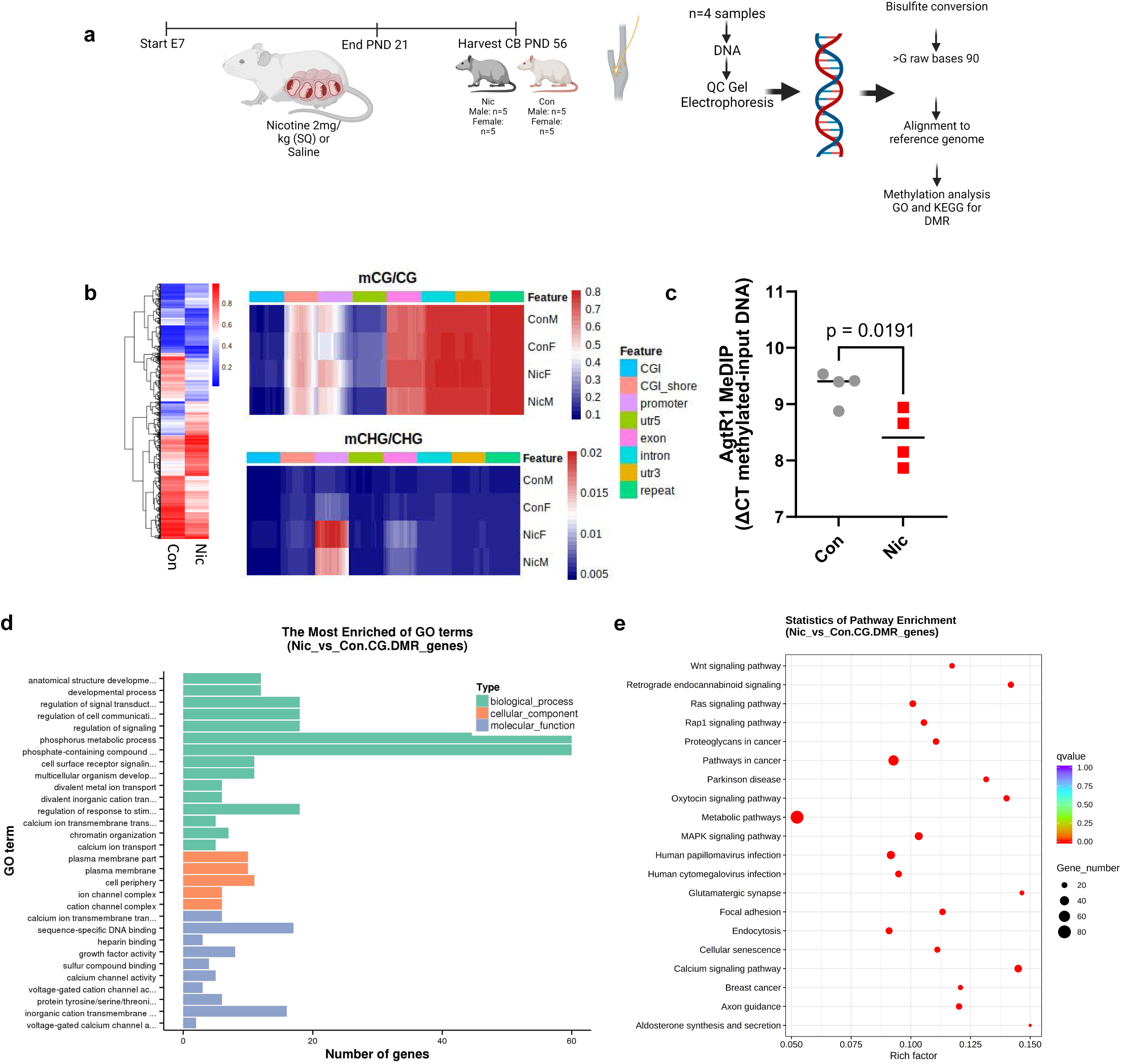
Perinatal nicotine exposure alters DNA methylation in the carotid body. **a)** DNA-methylation study design. Nicotine or saline was delivered to the mother on estrous day 7 (E7) to post-natal day (PND) 21. Bilateral carotid body (CB) samples were micro-dissected from perinatal nicotine or saline (control) exposed rats on PND 56. (n=5 rats per group-tissues were pooled to yield sufficient DNA for analysis). One male and one female group per condition. **b)** Heatmap displaying cytosine-guanine (C-G) and cytosine-arginine/threonine/guanine-guanine (CHG) methylation differences in carotid bodies between nicotine and saline-exposed rats. Mapping of methylation changes is confined to the CG and CHG regions as per the heat maps. **c)** Immunoprecipitation of methylated DNA compared to whole DNA demonstrates an increase of methylated AgtR1 DNA in the carotid bodies of nicotine exposed compared to control offspring. ΔCT was used to compare differences between groups (lower value equals greater amount of methylation). **d)** Enriched gene ontology (GO) terms for C-G methylated DNA between nicotine-exposed and control pups. **e)** Dot plot showing the gene number and enrichment factor of GO pathways between nicotine-exposed and control pups.

Proteins for AgtR1, TRPV1, and tyrosine hydroxylase (a standard marker of carotid body glomus cells) were all present in sections of carotid bodies harvested from perinatal nicotine-exposed offspring where AgtR1 is on glomus cells, and TRPV1 appears to oppose glomus cells on the post-synaptic terminals^38^ (Supplementary figure 1), However, our resolution is insufficient to determine the localization of glomus cells definitively. It is possible that these antibodies may suffer from this lack of specificity in rats. As such, our staining may not reflect the accurate location of AgtR1 in rat carotid body glomus cells. Nevertheless, given our RNAseq, qPCR, and DNA methylation data, along with others who have orthogonally demonstrated the presence of AgtR1 in glomus cells^38–41^, we are confident of the expression and upregulation of AgtR1 in the carotid bodies of perinatal nicotine exposed rats.

### Nicotine upregulates AgtR1 activity

To show that nicotine itself is responsible for AgtR1 upregulation, we used a cell culture model. PC12 cells have been routinely used as a surrogate carotid body glomus cell system^42–44^. Six hours of nicotine exposure significantly upregulated AgtR1a gene expression (Figure 3a). Differences in AgtR2 and TRPV1 were not demonstrated (Figure 3a). Differences in angiotensinogen, angiotensin converting enzyme, renin, and regulatory genes of EPAS1 (Hif2α) and tyrosine hydroxylase (TH-Figure 3a) were not demonstrated between conditions in relation to the housekeeping gene HPRT. DNA methylation of AgtR1 was confirmed with the DNA methylation pulldown assay (Figure 3b).

**Figure 3.**
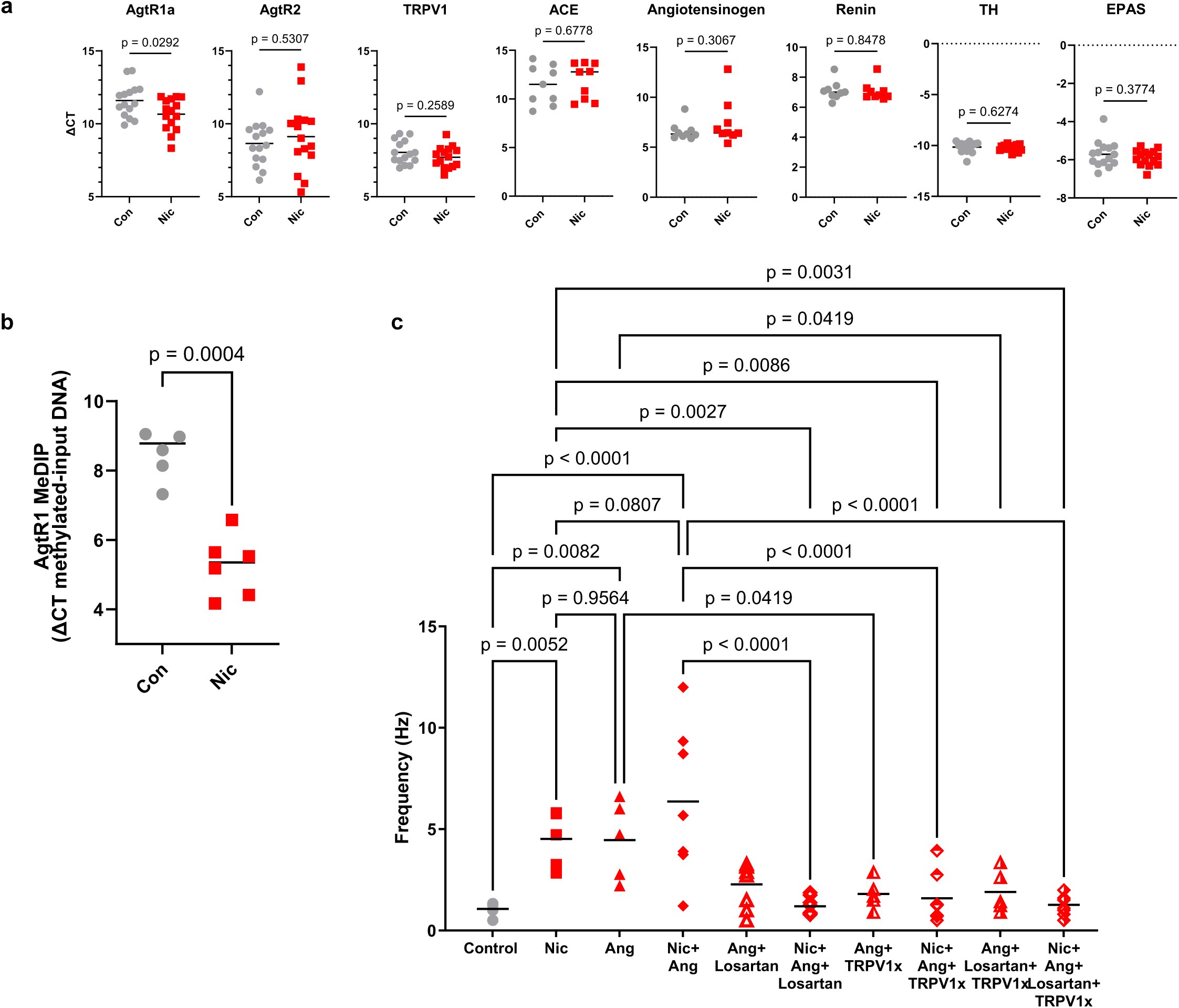
Nicotine increases AgtR1 pathway in cultured PC12 cells. **a)** PC12 cells (rat adrenal pheochromocytoma cells, resemble carotid body glomus cells) were cultured for 6 hours with and without 50µg/ml nicotine (similar concentration to a single dose of nicotine used *in vivo*), **a)** angiotensin II type 1a receptor (AgtR1a) was increased by nicotine (T_28_=2.299) The calculated increase of AgtR1a was 100.3% in comparison to control. Angiotensin II type 2 receptor (AgtR2) (T_28_=0.5307), transient receptor potential vanilloid 1 (TRPV1) (T_28_=0.2589), angiotensin converting enzyme (ACE, T_16_=0.4231), angiotensinogen (T_16_=1.056), renin (T_16_=0.1950), and regulatory genes of endothelial PAS domain-containing protein 1/ hypoxia inducible factor 2 alpha (EPAS1/Hif2α; T_28_=0.3774) and Tyrosine hydroxylase (TH-common PC12 and glomus cell marker, T_28_=0.6274) were unchanged. Data presented as ΔCT compared to Hypoxanthine-guanine phosphoribosyltransferase (HPRT) housekeeping gene. **b)** AgtR1 DNA was hypermethylated in PC12 cells following 6 hours of nicotine exposure. (T_10_=3.881). **c)** PC12 cells were cultured for 1 hour with media (control) nicotine (Nic, 50µg/ml), Angiotensin (Ang-50ng/ml), Nic + Ang, Ang+Losartan (AII1R blockade 3µM), Nic+Ang+Losartan, Ang+TRPV1 antagonist (TRPV1x, AMG9810 10µM), Nic+Ang+TRPV1x, Ang+Losartan+TRPV1x, and Nic+Ang+Losartan+TRPV1x and imaged for calcium flux using Incucyte calcium imaging with Calcium Orange (forms Calcium esther complex-non-ratiometric imaging). One-way ANOVA (F_9,54_=5.225, p<0.0001) Holm-Sidak post-hoc test, p-values inset in figure.

Using calcium imaging, we investigated how nicotine would augment glomus cell excitation. We show that with 1 hour of exposure to nicotine, PC12 cells increased their level of calcium spike frequency. The addition of angiotensin increased PC12 cell firing frequency to a level similar to nicotine. Importantly, nicotine combined with angiotensin did not further increase PC12 cell excitation. Further, losartan reduced calcium activity in PC12 cells when delivered to angiotensin or nicotine + angiotensin-stimulated cells concurrently. The reduction in calcium flux was also reduced with TRPV1 antagonist (AMG9810), and in combination with AMG9810 and Losartan (Figure 3c). In conclusion, nicotine has a stimulatory effect on PC12 and, quite likely, carotid body glomus cells to increase the expression and excitation of the angiotensin signaling pathway via AgtR1 and TRPV1.

### Angiotensin acutely increases arterial pressure

To assess acute carotid body reactivity to angiotensin, pups from perinatal nicotine-exposed or control mothers were anesthetized and given a bolus angiotensin injection into the jugular vein before and after acute carotid body denervation (Figure 4 a, b). In response to jugular vein injected angiotensin (circulates through the pulmonary circulation, then encounters the carotid bodies), arterial pressure was significantly elevated in nicotine-exposed offspring compared to controls (Figure 4c, Supplementary figure 4). In response to sodium cyanide, which blocks mitochondrial complex III and has been routinely used as a carotid body stimulant, there was no significant difference between groups (Figure 4d, Supplementary figure 4). Following carotid body denervation, arterial pressure was suppressed and did not respond to angiotensin or sodium cyanide as vigorously (Figure 4c, d, Supplementary figure 4). The reactivity to angiotensin and NaCN were attributed to the carotid bodies, however, given that boluses were delivered through the jugular vein, off-target effects cannot be ruled out.

**Figure 4.**
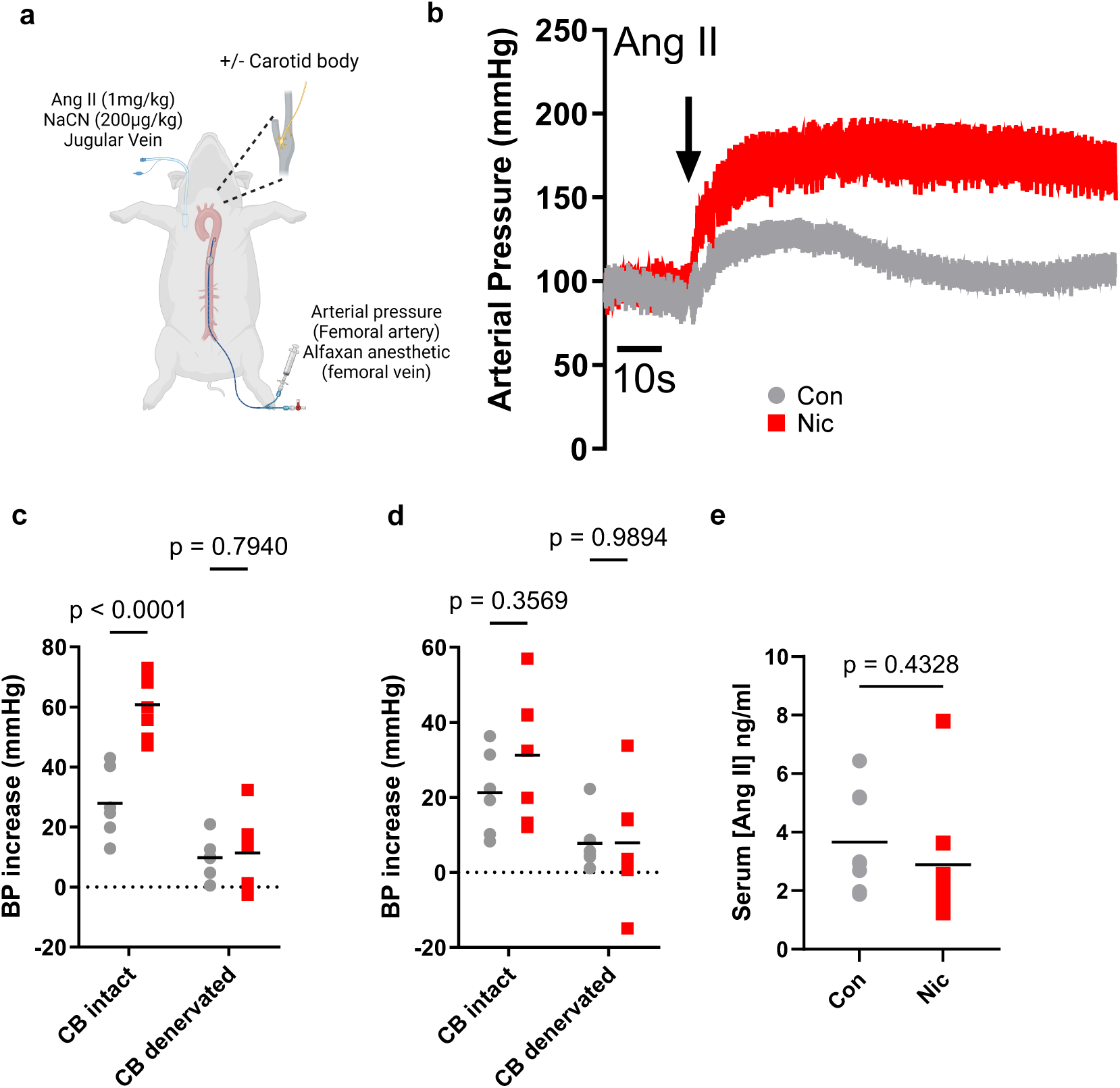
Carotid body reactivity to angiotensin is increased in perinatal nicotine-exposed pups compared to control with unaltered serum angiotensin. **a)** Diagram of *in vivo* experimental preparation-serum was isolated before injection of angiotensin or sodium cyanide (NaCN). **b)** A representative trace of arterial pressure of vehicle and perinatal nicotine-exposed pups on post-natal day (PND) 56 in response to a bolus jugular vein IV injection of angiotensin. **c)** The absolute increase in arterial pressure in response to angiotensin in vehicle (grey) and perinatal-nicotine exposed pups (red) before and following carotid body denervation F_1, 11_=11.34). **d)** The absolute increase in arterial pressure in response to NaCN in vehicle (grey) and perinatal-nicotine exposed pups (red) before and following carotid body denervation F_1, 11_=2.364). **e)** Serum was harvested before administration of bolus injections (T_14_=0.8077).

Importantly, serum harvested from these rats prior to any drug delivery demonstrated that circulating levels of angiotensin were not altered between groups (Figure 4e), similar to previous results^18^. It is possible that our analysis may be affected by background proteins in our sample, despite the ability of angiotensin II ELISAs to discern various forms of angiotensin and cleaved fragments^45^.

These data demonstrate a significant increase in carotid body excitability in response to angiotensin *in vivo* and suggest a hyperactive angiotensin signaling pathway within the carotid bodies of perinatal nicotine-exposed offspring.

### Carotid bodies are responsible for elevated blood pressure in perinatal nicotine-exposed offspring

We evaluated arterial pressure between quiescent conscious control and nicotine-exposed offspring using tail-cuff plethysmography on PND 56. Indeed, arterial pressure was higher in nicotine-exposed offspring (Figure 5a). To definitively demonstrate that the carotid bodies were primarily involved in perinatal nicotine-exposed hypertension, we implanted indwelling telemetry to measure real-time arterial pressure from the aorta and efferent sympathetic activity directed to the renal nerve between PND 56-60 (Figure 5b, Supplementary figure 5). We show that within the same animal, carotid body denervation drastically reduces arterial pressure and RSNA (Figure 5b, Supplementary figure 5). The reduction in arterial pressure with carotid body denervation was predominantly due to the suppression of systolic pressure, which significantly reduced pulse pressure (Supplementary figure 5). The reductions of RSNA as a result of carotid body denervation also coincide with a reduced respiratory rate, similar to other models of hypertension^19^. We attribute our changes in response to carotid body denervation to a reduction of RSNA as measured and demonstrated by others^19,20^. However, we did not assess the precise contribution of the carotid bodies to RSNA levels as previous^19^. Nevertheless, we note the association of RSNA to respiratory rate, which indicates that the reduction of respiratory rate following carotid body denervation would be tied to the fall in RSNA. These data demonstrate that carotid body basal activity and excitation contribute to the hypertensive state in perinatal nicotine-exposed offspring.

**Figure 5.**
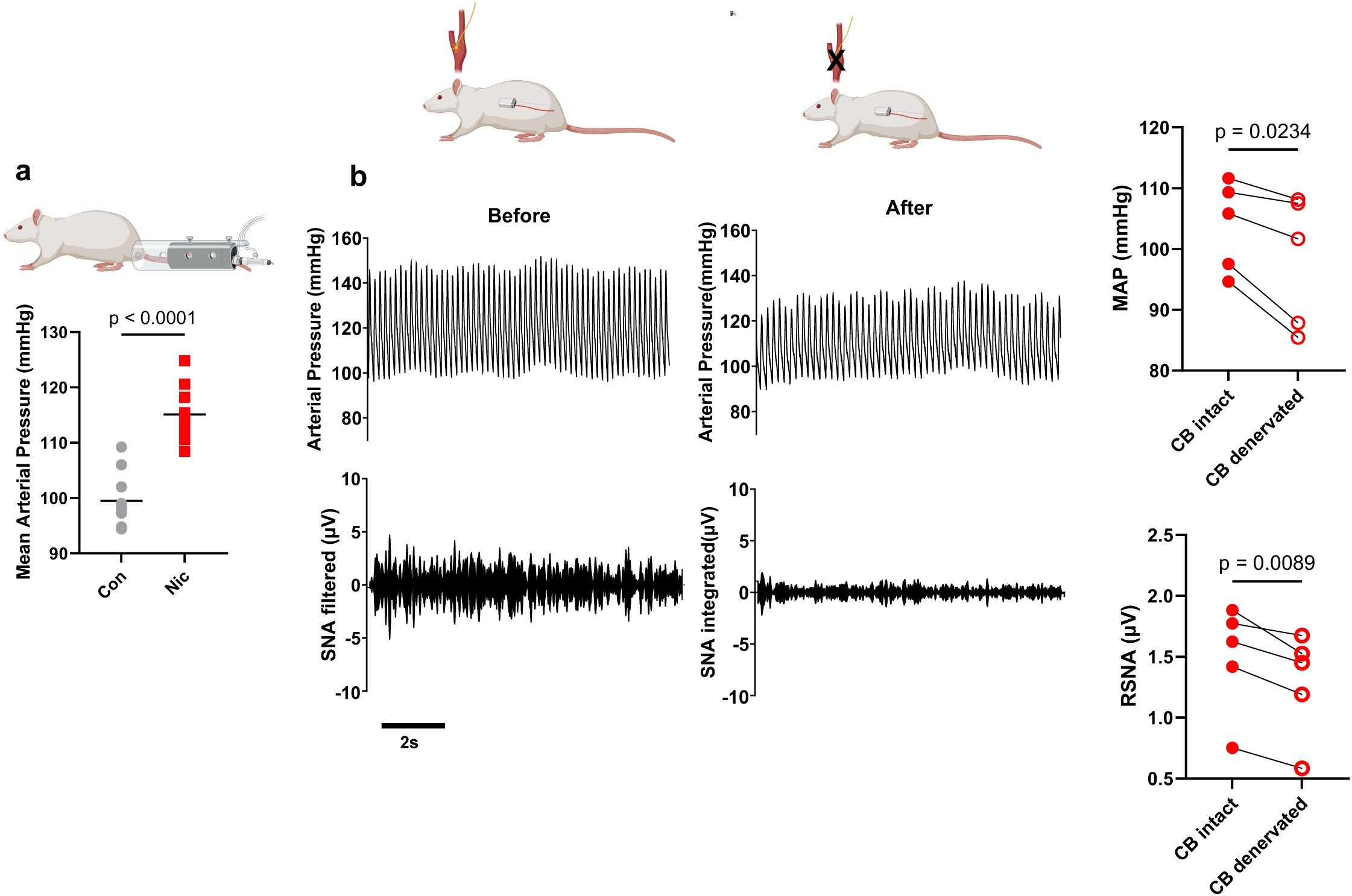
Perinatal nicotine-exposed rats have increased mean arterial pressure mediated by carotid body augmentation of sympathetic activity. **a**) Mean arterial pressure (MAP), as assessed by tail-cuff plethysmography, was augmented in nicotine-exposed offspring compared to controls (T_20_=7.964). N=11 per group, 2 independent experiments. **b**) Perinatal nicotine-exposed pups were implanted with radio telemetry to measure arterial pressure and renal sympathetic nerve activity (RSNA) on PND 56 as per Figure1a. Arterial pressure and renal sympathetic nerve activity was recorded for 7 days, and then carotid bodies were denervated and recordings recommenced for an additional 7 days. Raw traces of arterial pressure (AP) and efferent renal sympathetic nerve activity (vasoconstrictor outflow to kidneys) before carotid body denervation (Carotid body intact-BEFORE) and following carotid body denervation (Carotid body denervated-AFTER) in a representative perinatal nicotine exposed rat using telemetry. Summary statistics in n=5 perinatal nicotine rats comparing the absolute change of MAP and RSNA. MAP (T_4_=3.568), RSNA (T_4_=4.762).

## Discussion

Epigenetic changes in response to adverse maternal programming lead to cardiovascular sequelae in later life. Hypertension has been a consistent outcome associated with many adverse intra-uterine insults ranging from undernutrition^46,47^, overnutrition^47,48^, hypoxia^49,50^, and stress^51^. The risk of hypertension is thought to be significantly increased in perinatal nicotine-exposed offspring. The current data demonstrate that epigenetic changes in angiotensin signaling in the carotid bodies mediate hypertension in perinatal nicotine-exposed offspring.

We show, for the first time, that epigenetic modifications in response to adverse maternal programming in the form of nicotine exposure increase angiotensin, angiotensin II type 1a receptor transcripts, and angiotensin II type 1 receptor DNA intron methylation, demonstrating an increase in angiotensin signaling in the carotid bodies. These epigenetic changes *in vivo* are confirmed *in vitro* as we show that nicotine directly increases AgtR1 gene expression, AgtR1 DNA methylation, and AgtR1 stimulation in PC12, glomus-like cells, in culture.

Importantly, the difference in epigenetic changes to AgtR1 was predominantly confined to carotid bodies as the brain did not show an increase in gene expression or DNA methylation, whereas kidneys only showed increased DNA methylation between perinatal nicotine exposed and control pups. Angiotensinogen and angiotensin-converting enzyme gene expression was upregulated in the brains and kidneys, and renin was increased in the kidneys of perinatal nicotine exposed compared to control pups. In line with previous investigations, blood pressure reactivity to angiotensin is increased through a neurogenic mechanism involving carotid body-mediated excitation of efferent sympathetic activity. Finally, we showed, for the first time, that carotid body denervation reduced blood pressure in perinatal nicotine-exposed rats alongside efferent sympathetic activity.

Clinical data demonstrate that hypertension is likely in offspring that experienced intra-uterine/perinatal smoke exposure^2,7,52–56^; however, the age, sex, and potential for additional confounding lifestyle factors make it difficult to tease out this association, and measurements beyond adolescence have not been completed to date. Pre-clinical data using rodent models show that vascular reactivity to angiotensin via angiotensin II type 1 receptor gene expression, protein expression, and signaling increases in aortae and mesenteric arterioles following prenatal and perinatal nicotine exposure^14,15,18^. Accordingly, increased angiotensin II type 1 receptor activity, RNA, and DNA methylation^57,58^ have been consistent findings in response to perinatal nicotine-exposed offspring in vascular tissues. Our data show that additional sensory neural systems, which are significantly tied to hypertension^20^, are also augmented and induce hypertension in perinatal nicotine-exposed rats.

To date, scant data are available regarding the effects of nicotine exposure or perinatal nicotine exposure on carotid body activity and resulting autonomic reflex disturbances^23,59^. Carotid bodies are tasked with a multitude of homeostatic control mechanisms that regulate nutrient and oxygen delivery to tissues^21^. Their role in blood pressure regulation is highlighted by genetic^22^ and epigenetic^27^ modifications demonstrating the ability to increase basal activity and result in augmented blood pressure. Specifically, the spontaneously hypertensive rat (SHR) has augmented carotid body basal activity, which translates to alterations in breathing, blood pressure regulation, and glucose homeostasis^22^. Importantly, these carotid body changes appear to hinge on a significant difference in transcriptomic profile compared to their normotensive genetic counterpart, the Wistar-Kyoto (WKY) rat^22^. Further, and particularly salient to the current argument, carotid body denervation in SHR reduced arterial pressure and respiratory modulation, whereas carotid body denervation in WKY rats had no effect^19^. Importantly, evidence demonstrates that carotid body basal excitation can change via epigenetic modifications that suppress redox disinhibition^27^ to alter basal regulatory mechanisms such as breathing regulation.

A consistent finding is that intrauterine conditions with low oxygen^23^ or hyperleptinemia^60^ lead to augmented carotid body basal activity. Therefore, it is possible that deficiencies in the intrauterine environment augment arterial pressure in offspring due to the carotid body-regulated mechanism. Our data are the first to demonstrate that adverse maternal programming leads to carotid body epigenetic changes in AgtR1a and, importantly, the first to demonstrate that these epigenetic changes occur as a result of genetic upregulation via DNA methylation, which results in carotid body excitation. This carotid body excitation appears to be due to increased angiotensin signaling within the carotid bodies due to increases in carotid body micro signaling domains. Our *in vitro* experiments with PC12 cells demonstrate that epigenetic changes in carotid body angiotensin signaling directly result from nicotine exposure. Further, the reactivity of PC12 cells was regulated by nicotine to increase AgtR1 signaling. AgtR1 signaling in the carotid bodies has been tied to a TRPV1-mediated mechanism in response to repeated hypoxic bouts^38^. Given that losartan and TRPV1 blockade suppressed calcium signaling in PC12 cells, with no further reduction in calcium excitation with dual inhibition, we suggest that a similar mechanism is responsible for the augmentation of basal carotid body excitation. These data then fit with the increased carotid body reactivity to angiotensin in an anesthetized preparation and the consistent reduction in arterial pressure with carotid body denervation in conscious perinatal nicotine-exposed offspring.

## Conclusion

Hypertension resulting from perinatal nicotine exposure is due to epigenetic modifications within the carotid bodies. The increased AgtR1-TRPV1 signaling increases efferent sympathetic activity to augment arterial pressure. Future investigations should target neurogenic signaling mechanisms to alleviate hypertension due to adverse epigenetic programming.

### Perspectives

Carotid body denervation has shown clinical utility, as human clinical trials have shown that unilateral carotid body denervation results in a meaningful reduction in human blood pressure^61,62^. In fact, the reduction of systolic and diastolic arterial pressure with carotid body unilateral denervation^62,63^ appears to be similar to renal denervation^64,65^. Targeting the carotid bodies offers the advantage of suppressing the first step of reflex blood pressure regulation (i.e., sensory afferents^64,65^. Therefore, we suggest that a targeted suppression or reversal of angiotensin signaling in the carotid bodies and renin-angiotensinogen and angiotensin-converting enzyme in the kidneys would help alleviate increased blood pressure and cardiac afterload and limit unwanted side effects by standard drug therapies or invasive procedures.

## Acknowledgements and sources of funding

This project was supported by T32IP4707 of the Regents of the University of California Tobacco-Related Diseases Research Program (TRDRP). NJ is supported by NIH grant R21AI159221, R56AI175328, UCLA CTSI UL1TR001881-01 and, T32KT4708 of the Regents of the University of California TRDRP. WY is supported by NIH grant P50HD098593 and R01HD099924. VR is supported by NIH grant R01HL151769 and IR00737, T32IP5044, T32IR5048, and T32IR5365 of the Regents of the University of California TRDRP. The content is solely the responsibility of the authors and does not necessarily represent the official views of the National Institutes of Health or the UCOP TRDRP. Figures, 1a, 2a, 4a and 6a,b were completed with Biorender under a CC-BY-NC-ND license.

## Author contributions

F.Z. Z.W.& J.K. contributed to experimental design, conducted experiments and analyzed data. H.M, K.D. and J.L. contributed to experimental execution and data analysis. W.Y. and V.R. assisted with experimental design and analysis. N.J. designed the study, conducted experiments, analyzed data, and prepared the manuscript and figures. All authors agree on the manuscript.

