## Supplementary figures for "Epigenetic upregulation of carotid body angiotensin signaling increases blood pressure"

**Supplementary figure 1.**

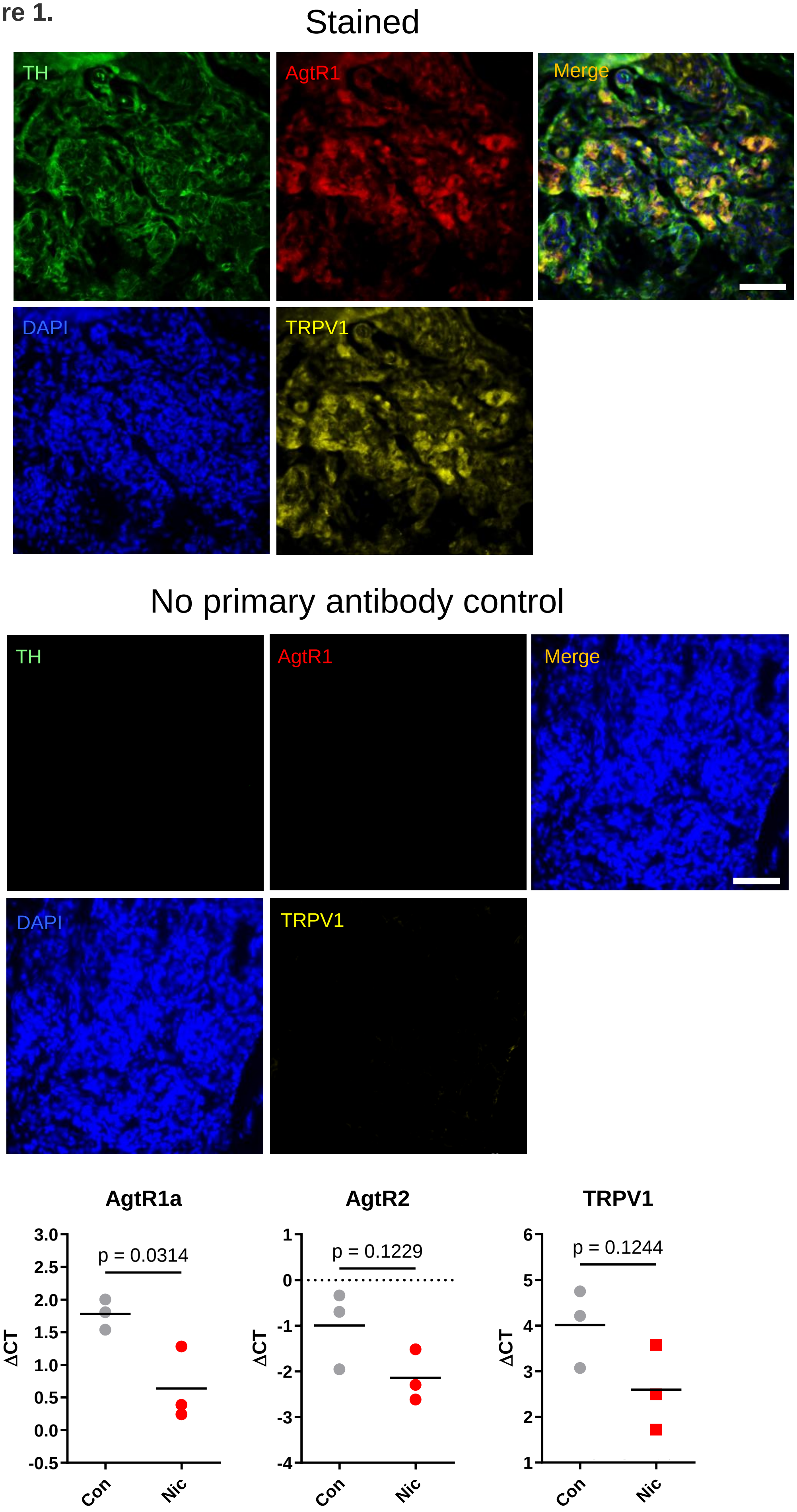


**Supplementary figure 1. Angiotensin II Type 1 receptor is colocalized with TRPV1 at in the carotid bodies.** Angiotensin II type 1 receptor (AgtR1) is colocalized to glomus cells (express tyrosine hydroxylase, TH) and are adjacent to transient receptor potential vanilloid 1 (TRPV1). Scale bar=20µm. Quantitative polymerase chain reaction (qPCR) of carotid bodies (n=3 per sample, n=3 samples per group) shows that AgtR1a is upregulated with perinatal nicotine exposure compared to saline control exposed rats. The calculated increase of AgtR1a was 56.45% in comparison to control. AgtR1a - $T_4$=2.792; AgtR2 - $T_4$=1.950; TRPV1-$T_4$=1.735.

**Supplementary figure 2.**

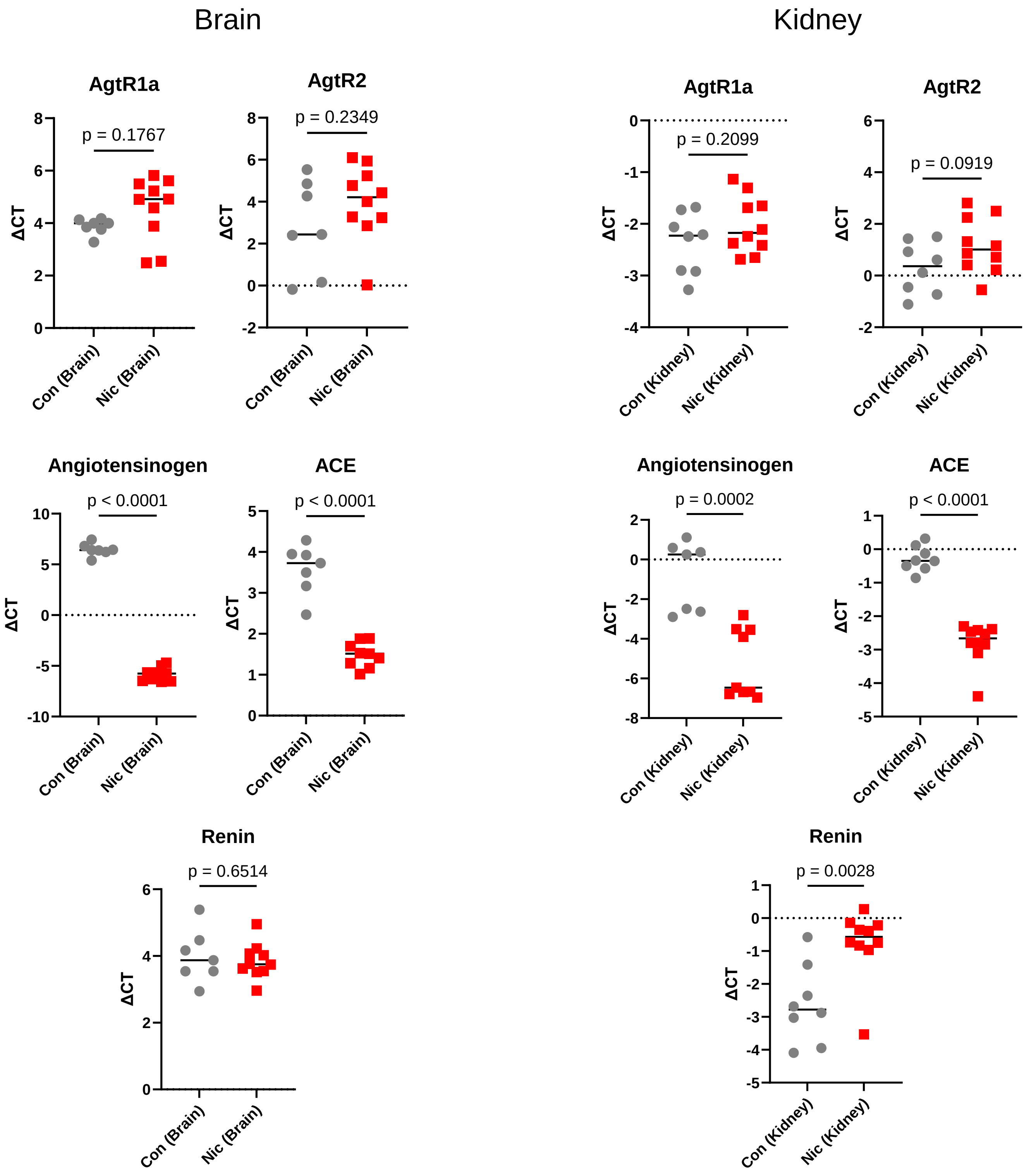


**Supplementary figure 2. Perinatal nicotine exposure increases gene expression for Angiotensin production in the brain and kidney.** Angiotensinogen and angiotensin converting enzyme are upregulated in brains and kidneys of perinatal nicotine exposed rats compared to control rats. Angiotensin receptors are unchanged between groups. Renin, in kidneys, is downregulated in perinatal nicotine exposed rats compared to control. Brain-AgtR1a- $T_{15}$=1.418, AgtR2- $T_{15}$=1.238, Angiotensinogen- $T_{14}$=37.16, angiotensin converting enzyme (ACE)- $T_{14}$=9.075, Renin- $T_{15}$=0.4610. Kidney- AgtR1a- $T_{16}$=1.306, AgtR2- $T_{16}$=1.793, Angiotensinogen- $T_{14}$=5.019, angiotensin converting enzyme (ACE)- $T_{16}$=10.12, Renin- $T_{16}$=3.529.

**Supplementary figure 3.**

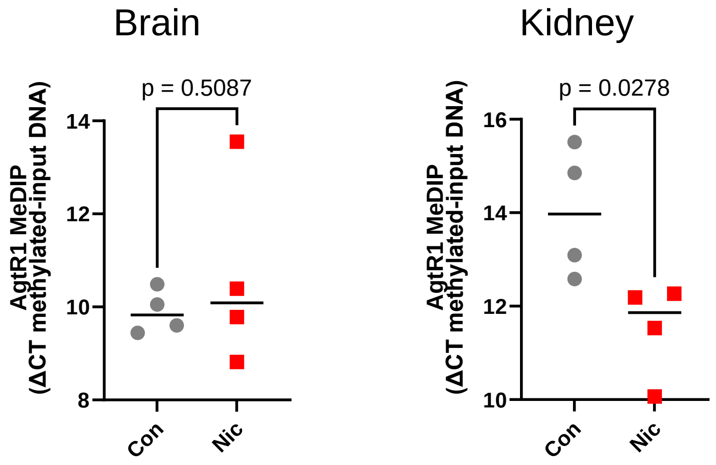


**Supplementary figure 3. DNA methylation of AgtR1 is confined to carotid bodies.** Immunoprecipitation of methylated DNA compared to whole DNA did not reveal differences between perinatal nicotine-exposed and control groups in the brains for AgtR1. AgtR1 methylated DNA was increased in kidneys of perinatal nicotine-exposed rats compared to controls. Brain- $T_6$=0.0946. Kidney- $T_6$=1.924.

### Supplementary figure 4.

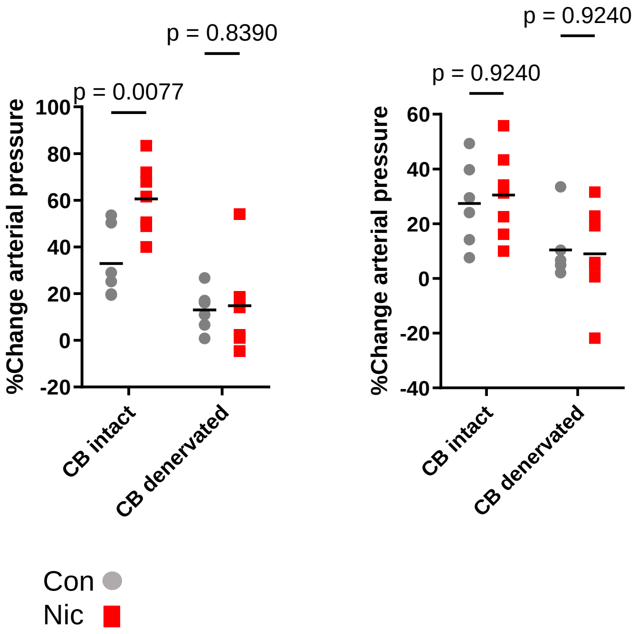


**Supplementary figure 4. Carotid body reactivity to angiotensin is increased in perinatal nicotine-exposed pups compared to control.** The percentage change in arterial pressure in response to angiotensin in vehicle (grey) and perinatal-nicotine exposed pups (red) before and following carotid body denervation $F_{1, 11}$=7.171). The percentage change in arterial pressure in response to NaCN in vehicle (grey) and perinatal-nicotine exposed pups (red) before and following carotid body denervation $F_{1, 11}$=0.6972). P-values for comparisons using Holm-Sidak post-hoc tests are inset in figure.

### Supplementary figure 5.

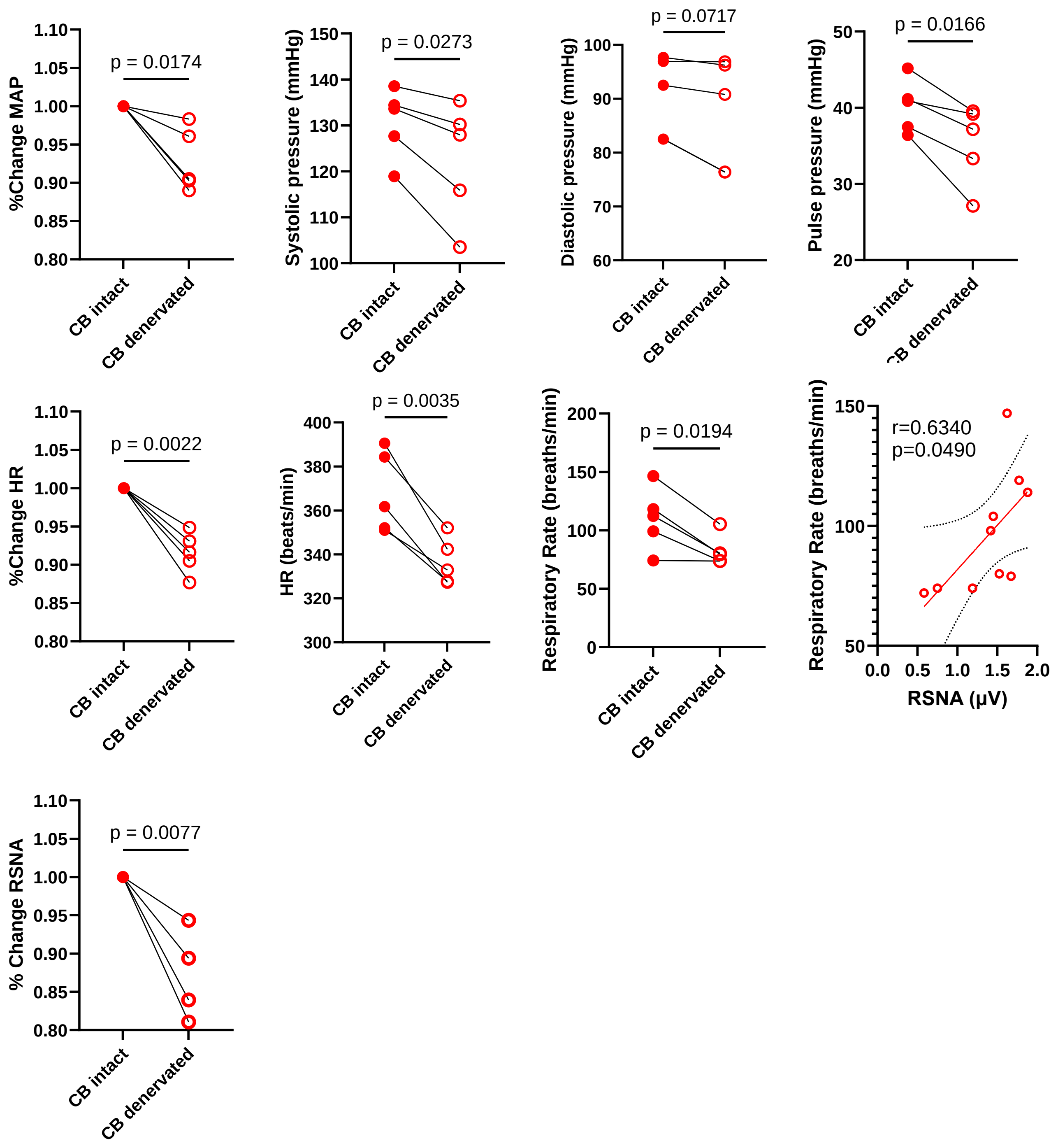


**Supplementary figure 5. Perinatal nicotine-exposed rats have increased mean arterial pressure mediated by carotid body augmentation of sympathetic activity.** Summary statistics in n=5 perinatal nicotine rats comparing the percent change of MAP($T_4$=3.908), Systolic pressure ($T_4$=3.399) and Diastolic pressure ($T_4$=2.433) Pulse pressure ($T_4$=3.966), %Change HR ($T_4$=6.971) and HR (beats/min, $T_4$=6.180), Respiratory rate (obtained from pressure waveform filtered at 300Hz low pass filter, $T_4$=3.783), Correlation of respiratory rate and RSNA, Pearson Correlation, and %Change RSNA($T_4$=4.956).
